# Improving Mannanase Production in *Bacillus subtilis* for Fibre Hydrolysis during Solid-State Fermentation of Palm Kernel Meal

**DOI:** 10.1101/2024.07.07.602432

**Authors:** Wei Li Ong, Zhi Li, Kian-Hong Ng, Kang Zhou

**Affiliations:** Wilmar Innovation Centre, Wilmar International Limited, 28 Biopolis Rd, Singapore 138568; Department of Chemical and Biomolecular Engineering, National University of Singapore

**Keywords:** *Bacillus subtilis*, Palm kernel meal, Mannanase, Solid-state fermentation

## Abstract

The primary challenge in utilizing palm kernel meal (PKM, an agricultural by-product) as non-ruminant livestock feed is its high fibre content, predominantly in the form of mannan. Microbial fermentation offers an economically favourable alternative to enzyme supplementation for breaking down fibre in lignocellulosic biomass. In a recent study, we have isolated and characterized an undomesticated strain (*Bacillus subtilis* F6) that is able to secrete mannanase. In this work, the mannanase production was substantially improved by optimizing multiple regulatory elements controlling the mannanase expression. Mannanase GmuG, sourced from *B. subtilis* F6 and verified for its hydrolytic activity on PKM fibre, was expressed using a replicative plasmid (pBE-S). The recombinant strain of *B. subtilis* F6 exhibited 1.9-fold increase in the mannanase activity during solid-state fermentation. Optimization of signal peptide and ribosome binding site further enhanced mannanase activity by 3.1-fold. Subsequently, promoter screening based on highly transcribed genes in *B. subtilis* F6 resulted in a significant 5.4-fold improvement in mannanase activity under the *nprE* promoter. The *nprE* promoter was further refined by eliminating specific transcription factor binding sites, enhancing the mannanase activity further by 1.8-fold. Notably, a substantial 35-40% reduction in PKM fibre content was observed after 30 h of fermentation using the recombinant strains. Lastly, the highest mannanase-producing strain was examined for scaled-up fermentation. The impacts of fermentation on fibre and protein contents, as well as the surface morphology of PKM, were analysed. The outcomes of this study offer an efficient method for robust mannanase expression in *B. subtilis* and its potential application in the biotransformation of PKM and other mannan-rich bioresources for improved feed utilization.

**Graphical abstract:** 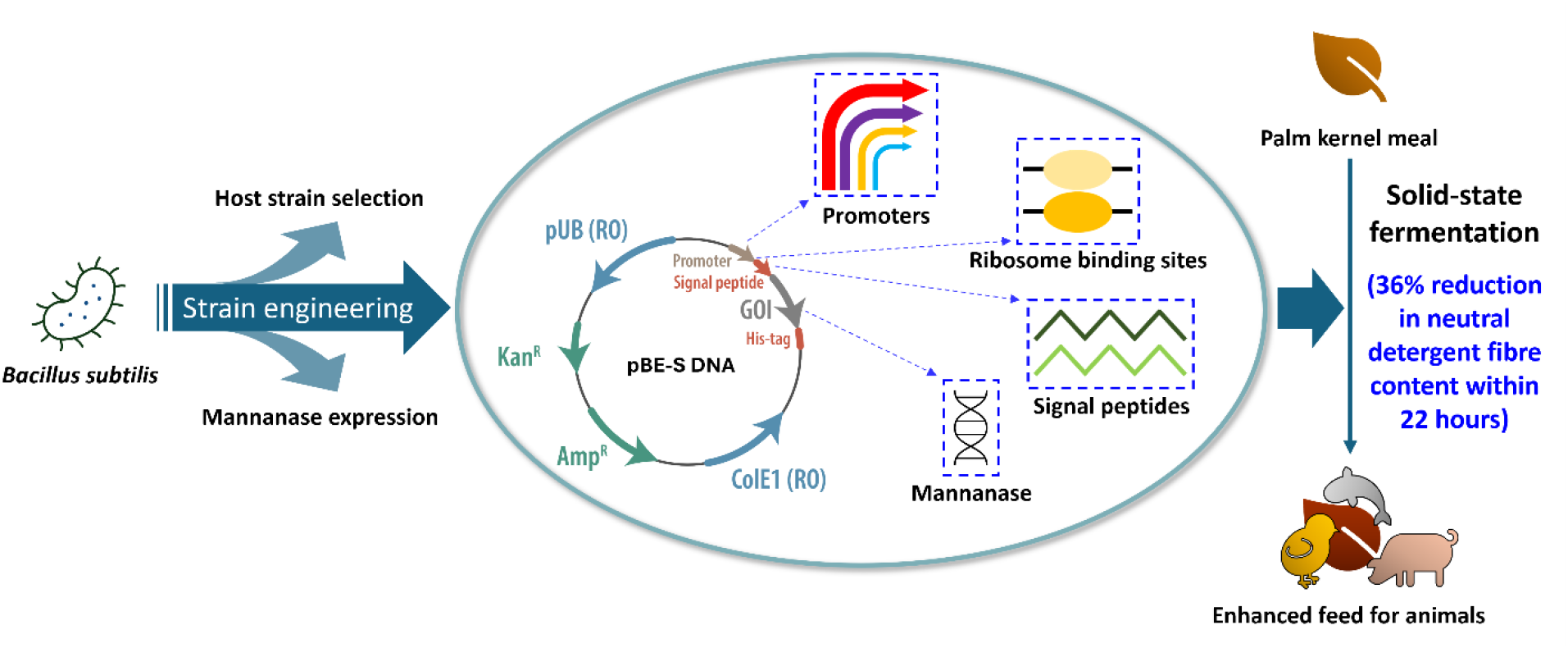

## 1. Introduction

Hemicellulose is the second most abundant polymer found in nature (Dawood & Ma, 2020). It is a vital component of plant cell walls. In grasses and hardwoods, xylan is the primary hemicellulose component, whereas in softwoods, plant fruits and seeds, hemicellulose predominantly exists in the form of mannan (Scheller & Ulvskov, 2010). Mannan is a polysaccharide containing D-mannose as the main monosaccharide constituent. The mannan in plants has both storage and structural functions (Capetti et al., 2023). Mannan mainly exists in four different forms: linear mannan, glucomannan, galactomannan, and galactoglucomannan. Due to the structural variations and heterogenous nature of mannan, its degradation may involve synergy among different enzymes including β-mannanase, β- mannosidase, β-glucosidase, α-galactosidase, and acetyl mannan esterase (Dawood & Ma, 2020).

As a major mannan-degrading enzyme, β-mannanase has attracted considerable attention from industry owing to its important applications across various sectors such as oil drilling, textiles, detergents, animal feed, pharmaceuticals, and the food industry (Capetti et al., 2023; Dawood & Ma, 2020). In animal feed industry, β-mannanase supplementation in high fibre diets has benefited the animals in many different ways. For instance, β-mannanase improved growth performance and reduced intestinal viscosity in broilers fed diets with varying levels of galactomannan (Latham et al., 2018). In addition, β-mannanase significantly improved weight gain, feed conversion ratio and specific growth rate in tilapia fed plant-based diet (Chen et al., 2016).

One of the mannan-rich feed ingredients that can potentially benefit from β-mannanase supplementation is palm kernel cake (PKC), which comes in two forms: solvent-extracted palm kernel meal (PKM) and mechanically pressed palm kernel expeller (PKE). Both are by- products of palm kernel oil extraction. Given that PKC is cheap, abundant, and produced year-around, it presents an attractive alternative to conventional feed ingredients such as corn and soybean meal. Although PKC has been satisfactorily used in the diet of ruminants, its utilisation efficiency in non-ruminant livestock feed is largely hindered by the high content of indigestible fibre, predominantly in the form of mannan (Sharmila et al., 2014).

Many studies have aimed to reduce the fibre content of PKC through direct enzyme supplementation or microbial fermentation. However, employing commercial enzymes for fibre hydrolysis proves economically challenging due to their high cost compared with the value of animal feed. Microbial fermentation emerges as a potentially cost-effective approach, enabling in situ enzyme production for fibre hydrolysis and biotransformation. Microbial degradation of mannan is primarily carried out by fungi such as *Aspergillus* species and *Rhizopus* sp., and Gram-positive bacteria including various *Bacillus* species. While microbial fermentation has been employed for fibre degradation in PKC, most studies involved prolonged fermentation time spanning several days, increasing production costs and contamination risks (**Table 1**). For instance, Marzuki et al. demonstrated a significant increase in crude protein content and decrease in crude fibre content after fermenting PKE with *Aspergillus niger* for 66 h (Marzuki et al., 2008). In another study, PKC fermented with *Bacillus amyloliquefacien* and *Trichoderma harzianum* resulted in reduced crude fibre content after a 7-day incubation period (Pasaribu et al., 2019). Similarly, better nutritional content was obtained in PKC after a 6-day fermentation using *Bacillus subtilis* (Mirnawati et al., 2019).

**Table 1.** Effect of solid-state fermentation on neutral detergent fibre (NDF) content of palm kernel cake (PKC)

| Fermenting microorganism (s) | Fermentation time | NDF before fermentation (%) | NDF after fermentation (%) | NDF hydrolysis (%) | Reference |
| --- | --- | --- | --- | --- | --- |
| *Bacillus subtilis* F6004np | 22 h | 78.9 | 42.6 | 36.4 | This work |
| *Bacillus subtilis* F6 | 22 h | 78.9 | 68.7 | 10.3 | This work |
| *Aspergillus niger* | 3 days | 73.64 | 66.57 | 7.07 | (Sitindaon et al., 2021) |
| *Bacillus amyloliquefaciens*,<br>*Trichoderma harzianum* | 7 days | 62.99 | 56.39 | 6.60 | (Pasaribu et al., 2019) |
| *Paenibacillus polymyxa* | 9 days | 82.29 | 71.70 | 10.59 | (Alshelmani et al., 2017) |
| *Rhizopus oryzae* | 7 days | 74.0 | 54.5 | 19.5 | (Ramin et al., 2010) |
| *Aspergillus niger* | 66 h | 79.0 | 50.3 | 28.7 | (Marzuki et al., 2008) |

Most studies have primarily examined the effects of solid-state fermentation on the nutritional values of PKE. Given that PKM is less preferred in livestock production due to its lower energy density, it stands to benefit more from biotransformation, leading to improved feed utilization. Therefore, PKM was selected as the subject for solid-state fermentation in our study. In a recent study, our group isolated a *B. subtilis* strain (F6) with a fast response time for mannanase production upon exposure to PKM, reducing the neutral detergent fibre (NDF) content of PKM by more than 10% in 24 h during solid-state fermentation (Ong et al., 2024). This work focuses on improving the mannanase production of the *B. subtilis* strain through genetic engineering to achieve greater fibre hydrolysis of PKM without extending fermentation time significantly. The mannanase gene was homologously expressed in a replicative plasmid.

To enhance the mannanase expression, plasmid construct was optimized by modifying the signal peptide (SP) and ribosomal binding site (RBS). Subsequently, promoter mining was conducted based on in-house transcriptome data by evaluating promoter candidates derived from genes highly expressed in strain F6 during PKM fermentation. The selected promoter P*nprE*, exhibiting the highest mannanase activity during fermentation, was further refined by eliminating specific transcription factor binding sites. Lastly, a time-course fermentation study was conducted to evaluate the performance of different recombinant strains of *B. subtilis* F6 in terms of mannanase production and PKM fibre hydrolysis. The highest mannanase- producing strain was subsequently examined for scaled-up fermentation. The effects of fermentation on fibre and protein contents, as well as the surface morphology of PKM, were assessed. As a result, by optimizing the different regulatory elements, the final production of mannanase was considerably increased by 60-fold compared to the wild-type strain in 30 h solid-state fermentation of PKM. Notably, a reduction of over 30% in NDF content was achieved after 22 h of fermentation using the recombinant strains (**Table 1**). This study is the first to report such efficient PKM fibre hydrolysis within a short fermentation time of one day using an engineered *B. subtilis* strain. This improvement not only prompted the potential application of engineered bacterial strain, but also implied the feasibility of the engineering strategies being applied to enhance the production of many other hydrolytic enzymes in *B. subtilis*.

## 2. Materials and methods

### 2.1 Strains, plasmids and cultivation conditions

Plasmids and bacterial strains used in this study were listed in **Tables 2** and **3**. *B. subtilis* strain F6 was isolated previously from exocarp of palm fruits (Ong et al., 2024).

*B. subtilis* strain CK7 was obtained from PT. Wilmar Bernih Indonesia (Virginia et al., 2018). *B. subtilis* strain 1A1 was obtained from Bacillus Genetic Stock Center (BGSC). *B. subtilis* strain RIK1285 and *B. subtilis*/ *E. coli* shuttle vector pBE-S DNA were purchased from Takara Bio. *E. coli* and *B. subtilis* strains were cultured at 37 °C with 220 rpm orbital shaking in lysogeny broth (LB) medium. Appropriate antibiotics were supplemented to the medium when required: 100 μg/mL ampicillin for *E. coli* and 10 μg/mL kanamycin for *B. subtilis*.

**Table 2.**
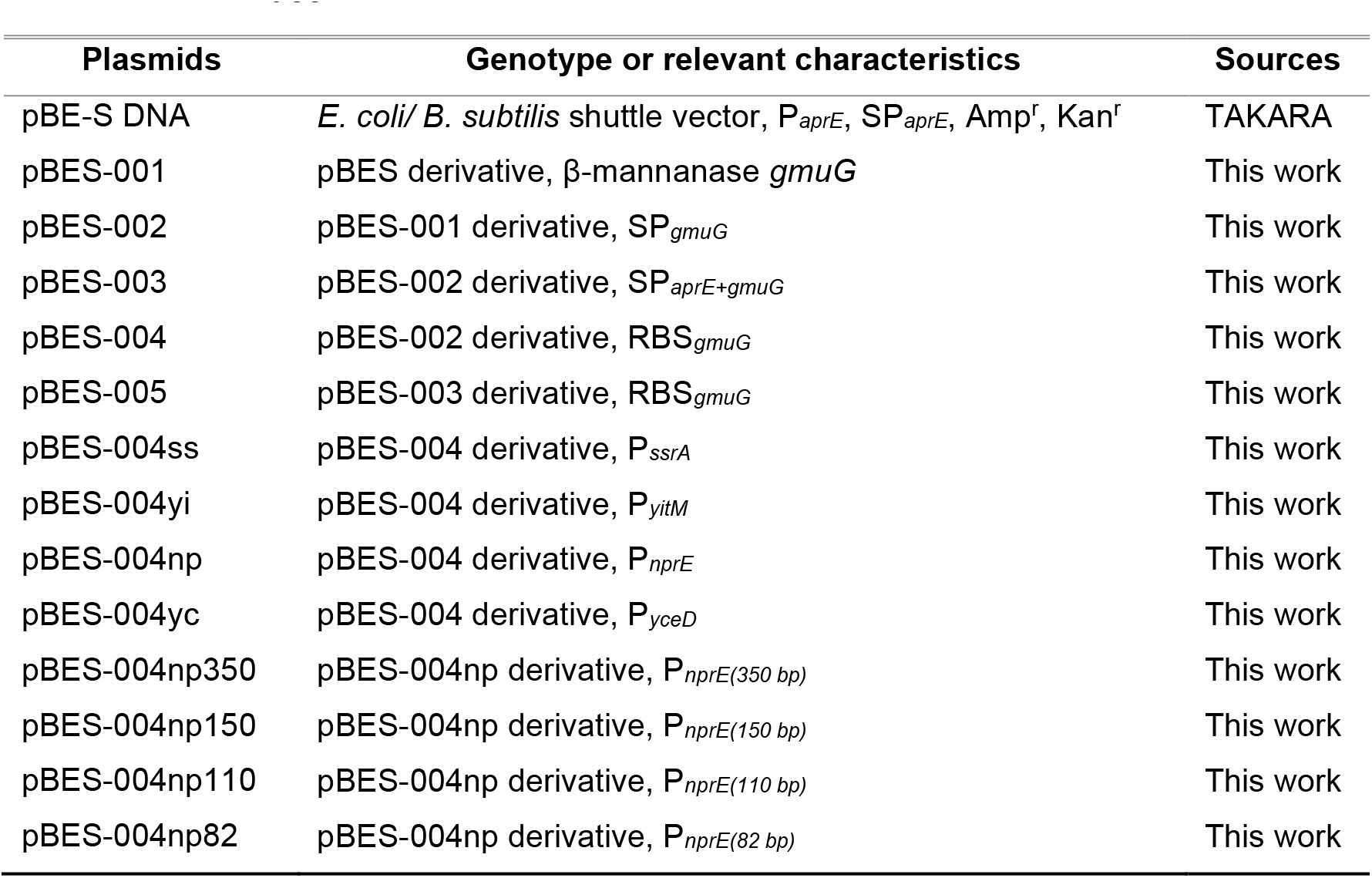
Plasmids used in this study.

| Plasmids | Genotype or relevant characteristics | Sources |
| --- | --- | --- |
| pBE-S DNA | *E. coli*/ *B. subtilis* shuttle vector, P<sub>aprE</sub>, SP<sub>aprE</sub>, Amp<sup>r</sup>, Kan<sup>r</sup> | TAKARA |
| pBES-001 | pBES derivative, β-mannanase *gmuG* | This work |
| pBES-002 | pBES-001 derivative, SP<sub>*gmuG*</sub> | This work |
| pBES-003 | pBES-002 derivative, SP<sub>*aprE+gmuG*</sub> | This work |
| pBES-004 | pBES-002 derivative, RBS<sub>*gmuG*</sub> | This work |
| pBES-005 | pBES-003 derivative, RBS<sub>*gmuG*</sub> | This work |
| pBES-004ss | pBES-004 derivative, P<sub>*ssrA*</sub> | This work |
| pBES-004yi | pBES-004 derivative, P<sub>*yitM*</sub> | This work |
| pBES-004np | pBES-004 derivative, P<sub>*nprE*</sub> | This work |
| pBES-004yc | pBES-004 derivative, P<sub>*yceD*</sub> | This work |
| pBES-004np350 | pBES-004np derivative, P<sub>*nprE*(350 bp)</sub> | This work |
| pBES-004np150 | pBES-004np derivative, P<sub>*nprE*(150 bp)</sub> | This work |
| pBES-004np110 | pBES-004np derivative, P<sub>*nprE*(110 bp)</sub> | This work |
| pBES-004np82 | pBES-004np derivative, P<sub>*nprE*(82 bp)</sub> | This work |

**Table 3.**
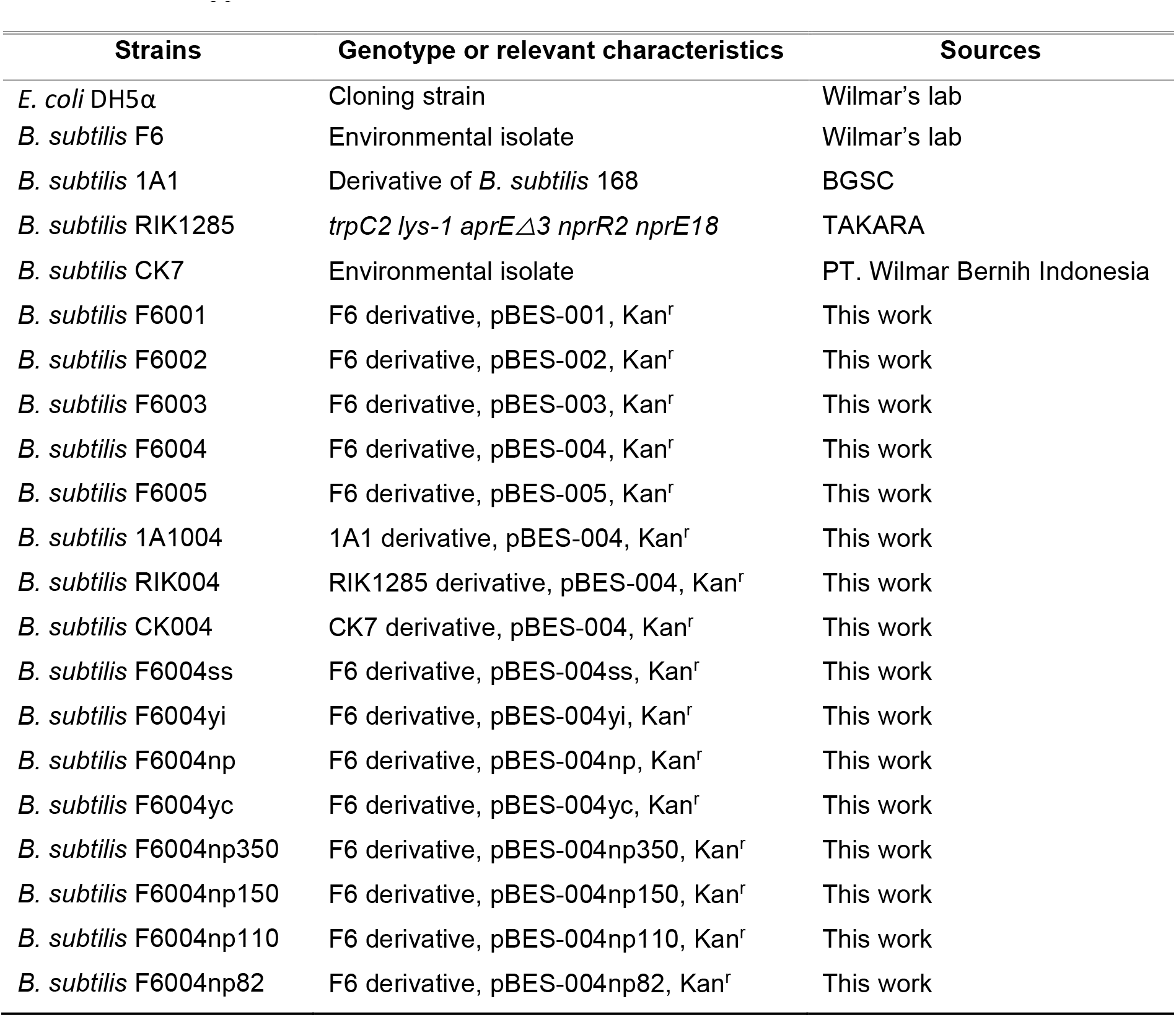
Strains used in this study.

| Strains | Genotype or relevant characteristics | Sources |
| --- | --- | --- |
| *E. coli* DH5α | Cloning strain | Wilmar's lab |
| *B. subtilis* F6 | Environmental isolate | Wilmar's lab |
| *B. subtilis* 1A1 | Derivative of *B. subtilis* 168 | BGSC |
| *B. subtilis* RIK1285 | *trpC2 lys-1 aprE*Δ3 *nprR2 nprE18* | TAKARA |
| *B. subtilis* CK7 | Environmental isolate | PT. Wilmar Bernih Indonesia |
| *B. subtilis* F6001 | F6 derivative, pBES-001, Kan<sup>r</sup> | This work |
| *B. subtilis* F6002 | F6 derivative, pBES-002, Kan<sup>r</sup> | This work |
| *B. subtilis* F6003 | F6 derivative, pBES-003, Kan<sup>r</sup> | This work |
| *B. subtilis* F6004 | F6 derivative, pBES-004, Kan<sup>r</sup> | This work |
| *B. subtilis* F6005 | F6 derivative, pBES-005, Kan<sup>r</sup> | This work |
| *B. subtilis* 1A1004 | 1A1 derivative, pBES-004, Kan<sup>r</sup> | This work |
| *B. subtilis* RIK004 | RIK1285 derivative, pBES-004, Kan<sup>r</sup> | This work |
| *B. subtilis* CK004 | CK7 derivative, pBES-004, Kan<sup>r</sup> | This work |
| *B. subtilis* F6004ss | F6 derivative, pBES-004ss, Kan<sup>r</sup> | This work |
| *B. subtilis* F6004yi | F6 derivative, pBES-004yi, Kan<sup>r</sup> | This work |
| *B. subtilis* F6004np | F6 derivative, pBES-004np, Kan<sup>r</sup> | This work |
| *B. subtilis* F6004yc | F6 derivative, pBES-004yc, Kan<sup>r</sup> | This work |
| *B. subtilis* F6004np350 | F6 derivative, pBES-004np350, Kan<sup>r</sup> | This work |
| *B. subtilis* F6004np150 | F6 derivative, pBES-004np150, Kan<sup>r</sup> | This work |
| *B. subtilis* F6004np110 | F6 derivative, pBES-004np110, Kan<sup>r</sup> | This work |
| *B. subtilis* F6004np82 | F6 derivative, pBES-004np82, Kan<sup>r</sup> | This work |

### 2.2 Genetic manipulation

The mannanase gene *gmuG* (without the native signal peptide) derived from *B. subtilis* strain F6 was cloned into the vector pBE-S DNA (Takara Bio). The genome of *B. subtilis* strain F6 also served as the template to amplify signal peptide SP*gmuG*, ribosome binding site RBS*gmuG* and promoter fragments including P*ssrA*, P*yitM*, P*yceD* and P*nprE*. The plasmid was first constructed in *E. coli* DH5α and then introduced into *B. subtilis* by using electroporation.

### 2.3 Palm kernel meal solid-state fermentation (SSF)

PKM was obtained from Wilmar palm kernel solvent extraction plant located in East Java, Indonesia. PKM was sterilized at 121°C for 15-20 min before fermentation. *B. subtilis* cells were precultured overnight in LB medium, and a cell pellet containing 6×10^8^ CFU was resuspended in 1.5 ml of deionized water. This suspension was then used to inoculate 1 g of PKM in a 50 ml tube, which was incubated at 37 °C for 22-24 h. After fermentation, 10 ml of deionized water was added to the fermented PKM, briefly vortexed at high speed, and centrifuged to obtain the cell-free supernatant for mannanase activity testing and soluble sugar analysis. Alternatively, the fermented PKM was dried and stored at room temperature for subsequent fibre and protein analysis.

### 2.4 Determination of mannanase activity

Mannanase activity was assayed by using a 0.5% (w/v) solution of locust bean gum (LBG) galactomannan in 100 mM sodium acetate buffer pH 5.0 (950 µl) to which 50 µl of the appropriately diluted enzyme solution was added. The reducing sugars released in 20 min at 55°C were measured as mannose equivalents by the dinitrosalicylic acid (DNS) method described by Miller (Miller, 1959) with minor modifications. Briefly, 300 µl of the reaction mixture was reacted with 900 µl of DNS reagent solution at 100°C for 5 min followed by absorbance measurement at 540 nm. 500 ml of DNS reagent solution was prepared with 5 g DNS, 5 g sodium hydroxide pellet, and 100 g Rochelle salt dissolved in water. One unit of mannanase activity is defined as the amount of enzyme producing 1 µmol of mannose equivalents per min under the given conditions.

### 2.5 SDS-PAGE and western bot analysis

LB liquid culture and resuspended solid culture were centrifuged to obtain supernatant for protein analysis using sodium dodecyl sulfate polyacrylamide gel electrophoresis (SDS-PAGE). Samples were resolved on Mini-PROTEAN TGX Gels (4-20%) (Biorad, USA) followed by staining with quick Coomassie stain (Protein Ark, UK). Prestained protein ladder (Bio-Rad, USA) was applied to determine the molecular weights of the proteins with prominent bands. For western blot analysis, proteins separated by SDS-PAGE were transferred to nitrocellulose or PVDF membranes. Membranes were blocked with 5% milk in PBST overnight at 4°C, then incubated with anti-GmuG primary antibody for 1-2 h at room temperature. After washing, membranes were incubated with HRP-linked anti-rabbit IgG secondary antibody (Cell Signaling Technology, USA) for 1-2 h at room temperature. Signals were detected using Pierce ECL western blotting substrate (Thermo Scientific, USA) and imaged with the ChemiDoc Imaging System (Bio-Rad, USA).

### 2.6 High performance liquid chromatography (HPLC) for sugar analysis

The sugar quantification was done using HPLC with a refractive index detector (RID). The chromatography was performed with Agilent 1260 Infinity II, installed with Agilent Hi-Plex Na guard column (7.7 x 50 mm). Separation of oligosaccharides was achieved using Agilent Hi-Plex Na analytical column (7.7 x 300 mm) in 45 min with an isocratic flow of 0.3 ml/ min using 100% Milli-Q water. Samples were injected using an auto- sampler and the injection volume was 5 μl. Column temperature was 85 °C. The detector signal was expressed as nano Refractive Index Units (nRIU). The standards (mannohexaose, mannopentaose, mannotetraose, mannotriose and mannobiose) were purchased from Megazyme (Ireland).

### 2.7 RNA extraction and transcriptome analysis

The overnight LB culture of *B. subtilis* F6 was centrifuged to obtain a cell pellet containing 12x10^8^ CFU, which was then resuspended in 3 ml of sterile water. This suspension was added to 2 g of autoclaved PKM in a 50 ml tube for SSF at 37°C for 6 h. After fermentation, the fermented PKM was mixed with 8 ml phosphate-buffered saline (PBS) followed by filtration through miracloth (Sigma) to remove PKM debris and collect bacterial cells for RNA extraction. Total RNA was isolated using RNeasy Protect Bacteria Mini Kit (Qiagen, USA) including on-column DNase digestion. Quantifications and integrity were checked using RNA ScreenTape Assay with Agilent 4200 TapeStation System (Agilent). All samples had an RNA integrity number (RIN) > 9. Subsequently, whole transcriptome sequencing was performed in order to examine the gene expression profiles (Macrogen, South Korea). The library kit and type of sequencer were TruSeq Stranded Total RNA (NEB Microbe) and NovaSeq, respectively. Read mapping was performed with *B. subtilis* BSn5 as the reference strain. Transcript expression level was quantified in term of RPKM (Reads Per Kilobase of transcript per Million mapped reads) values.

### 2.8 Determination of neutral detergent fibre (NDF) content

0.5 g of the dried PKM sample was used for NDF analysis according to the protocol based on AOAC 2003:04/ ISO 16472:2006. In brief, the sample was treated with sodium sulphite and neutral detergent solution using the default program in fibre analyzer (Fibertec 8000, FOSS). 1 L of neutral detergent solution with a final pH between 6.95 to 7.05 was prepared with 18.61 g of EDTA (disodium ethylene diamine tetraacetate), 6.81 g of sodium borate decahydrate, 30 g sodium lauryl sulphate, 10 ml of triethylene glycol and 4.56 g disodium hydrogen phosphate dissolved in water.

### 2.9 Scaled-up PKM solid-state fermentation

The scaled-up fermentation was performed by multi-step blending. Firstly, five individual tubes of 1 g PKM with an initial inoculum of 6 x 10^8^ CFU each was allowed to ferment for 6 h. Next, the pre-fermented mixture of PKM and *B. subtilis* cells was used as the starter to inoculate another 15 g of PKM in a 300 ml flask. After 18 h of incubation, 20 g of fermented PKM was mixed with 40 g of PKM in a 1 L flask to undergo 24 h fermentation to get the final fermented product. In each blending step, 1.5 ml water was added for every 1 g of fresh PKM introduced into the system. The fermentation was performed at 37 °C. The fermented PKM was dried and stored at room temperature until analysis of surface morphology, fibre and protein contents.

### 2.10 Determination of protein content

1 g of the dried PKM sample was used for analysis of Kjeldahl nitrogen using the FOSS Kjeltec System. The sample was subjected to block digestion followed by steam distillation. During digestion, the nitrogen or protein in the sample was converted to ammonium sulphate. Next, the ammonium was converted into ammonia by using an alkali (NaOH). The ammonia was steam distilled into a receiver flask containing boric acid. Lastly, titration with standard acid solution was performed using colorimetric end- point detection. To convert nitrogen content into protein content, a conversion factor of 6.25 was used.

### 2.11 Analysis of PKM surface morphology using scanning electron microscopy

The dried PKM samples were observed under a field emission scanning electron microscopy (FE-SEM) JSM-7610F Plus (Jeol, Japan) . Briefly, the PKM sample was placed onto a 10 mm copper sample stub for the examination of the sample surface morphology under FE-SEM at 450 x magnification.

## 3. Results and discussion

### 3.1 Enhanced extracellular production of β-mannanase by changing signal peptide and ribosomal binding site

The replicative plasmid pBES-001 was constructed to express the β-mannanase *gmuG* gene from *B. subtilis* F6 under the P*aprE* promoter with the SP*aprE* signal peptide (**Figure 1A**). Upon transformation of *B. subtilis* F6 with plasmid pBES-001, the engineered strain F6001 showed a 9- and 1.9-fold increase in its mannanase activity compared to the wild-type strain under overnight LB culture and PKM fermentation conditions, respectively (**Figure 1B, C**). The secretory expressions of recombinant GmuG protein under both growth conditions were verified by western blot (**Figure 1D, E**). It was found that the native strain of *B. subtilis* F6 produced very low level of mannanase activity in LB medium compared to growing in the presence of PKM. Mannanase overexpression using the multicopy plasmid substantially improved the mannanase activity of the strain F6 in LB medium. Nonetheless, the improvement of mannanase activity during PKM fermentation was limited, necessitating the optimization of plasmid construct for higher mannanase expression.

**Figure 1.**
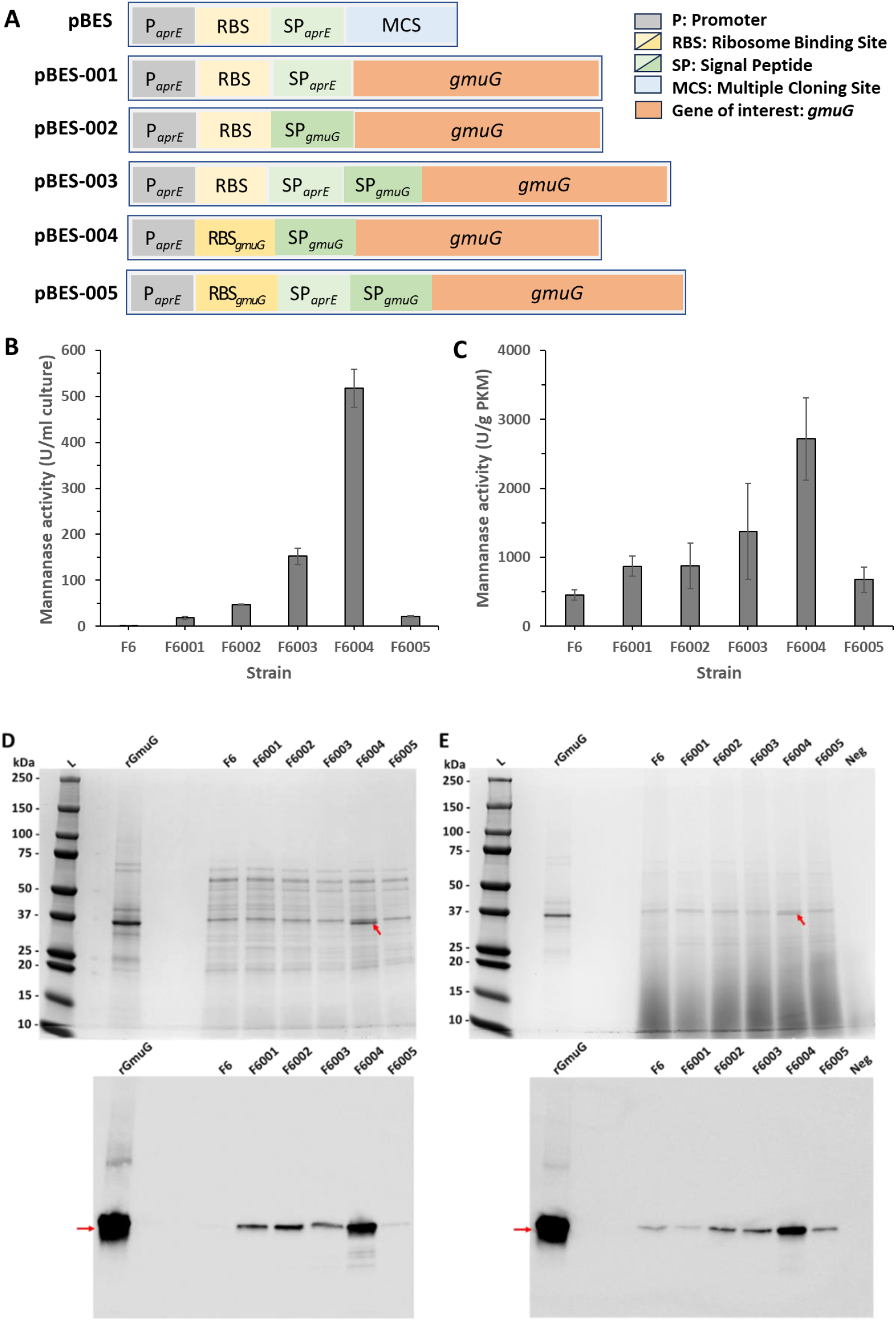
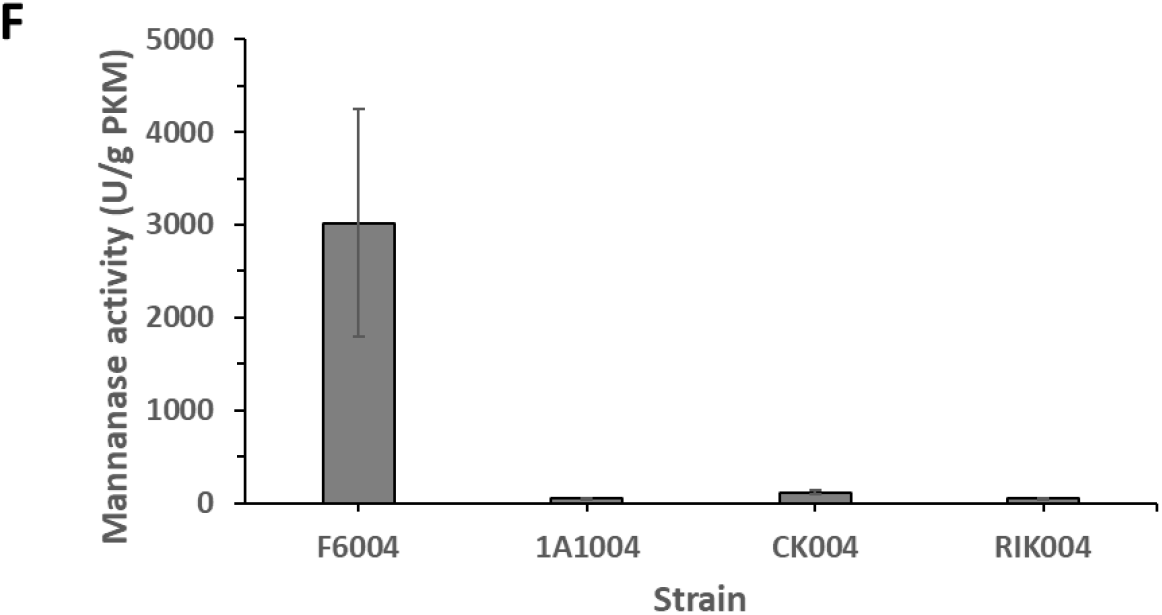
Optimization of signal peptide and ribosome binding site for GmuG overexpression. (A) Schematic diagram showing the different pBES plasmid constructs. P: promoter; RBS: ribosome binding site; SP: signal peptide; MCS: multiple cloning site. (B, C) Mannanase activity in (B) overnight LB culture and (C) PKM solid-state fermentation. Data are the mean of triplicates, and bars indicate standard deviation of triplicates. (D, E) SDS-PAGE (top) and western blot (bottom) analysis of GmuG expression in (D) overnight LB culture and (E) PKM solid-state fermentation. The band corresponding to intact GmuG was marked. L: protein ladder; rGmuG: purified recombinant GmuG protein as positive control for western blot; Neg: PKM negative control. (F) Mannanase activity of different recombinant strains of *B. subtilis* containing the optimized plasmid pBES-004 in PKM solid-state fermentation. Data are the mean of triplicates, and bars indicate standard deviation of triplicates.

In *B. subtilis*, the export of protein is generally accomplished by the Sec-type secretion pathway (Fu et al., 2018). The N-terminal sequence of a secreted protein carries a specific secretion signal known as signal peptide. Apart from being required for the targeting and membrane translocation by the respective protein translocases, signal peptides also have additional influences on the biosynthesis, the folding kinetics, and the stability of the respective target proteins (Freudl, 2018). Therefore, for efficient secretory production of recombinant proteins, the choice of the signal peptide is an important factor to consider. Here, the *gmuG* signal peptide (SP*gmuG*) derived from *B. subtilis* F6 was chosen to replace the *aprE* signal peptide (SP*aprE*) of plasmid pBES-001, generating the plasmid pBES-002 (**Figure 1A**). Upon transformation of *B. subtilis* F6 with plasmid pBES-002, the engineered strain F6002 exhibited a 2.6-fold increase in mannanase activity in overnight LB culture compared to strain F6001 (**Figure 1B**). However, under PKM fermentation conditions, strain F6002 did not demonstrate any improvement in activity compared to strain F6001 (**Figure 1C**).

Individual signal peptide has their own merits and disadvantages in terms of the effects on signal peptide processing, levels of precursor or mature protein accumulation, and production of properly folded protein (Zhang, 2015). Hence, the combination of two signal peptides may offer some advantages over the traditional Sec-dependent signal peptides. Here, the SP*gmuG* of plasmid pBES-002 was replaced with a synthetic dual signal peptide combining SP*aprE* and SP*gmuG*, yielding the plasmid pBES-003 (**Figure 1A**). Upon transformation of *B. subtilis* F6 with plasmid pBES-003, the engineered strain F6003 exhibited a 3.2-fold increase in mannanase activity in LB culture compared to strain F6002 (**Figure 1B**). In PKM fermentation, the strain F6003 showed a 1.6-fold increase in mannanase activity compared to strain F6002 (**Figure 1C**). The use of dual signal peptide was previously investigated for the secretion of maltose binding protein (MBP) in *E. coli* (Zhang, 2015).

RBS sequences have been reported to play an important role in initiating translation and controlling protein expression levels (Shi et al., 2020). Based on the two plasmids pBES-002 and pBES-003, the 19-bp sequence ‘CAAAAGGAGAGGGACGCGT’ containing the vector-derived RBS ‘AGGAG’ was replaced by a 20-bp sequence ‘AATGAATGGGGGAGTTGCAT’ containing the native ribosome binding site of *gmuG* gene (RBS*gmuG*), generating new constructs pBES-004 and pBES-005 (**Figure 1A**), respectively. The two plasmids were used to transform *B. subtilis* F6, creating the two corresponding plasmid-containing strains F6004 and F6005, respectively. Among the recombinant strains (F6001 to F6005), the highest mannanase activity was observed with strain F6004, in which both the RBS and SP were derived from the *gmuG* gene (**Figure 1B, C**). The activity was improved significantly by 11- and 3-fold compared to the strain F6002 under LB culture and PKM fermentation conditions, respectively. As shown in SDS-PAGE and western blot (**Figure 1D, E**), strain F6004 also showed the highest mannanase expression under both growth conditions. The improved enzyme activity could be attributed to the changes in the efficiency of translation initiation. Previous research suggests that interactions between the spacer sequence (located between the RBS and the start codon of the target gene) and the 5’ region of the target gene (which encodes the signal peptide for Sec-secreted proteins) can influence translation initiation (Volkenborn et al., 2020). Both the 5’ UTR and the 5’ end of the coding sequence can affect mRNA secondary structures, potentially masking the RBS and preventing ribosome binding.

*B. subtilis* produces various extracellular proteases that can degrade target proteins (Liu et al., 2018), so protease-deficient strains like RIK1285 were used to improve protein expression (Murayama et al., 2004). The optimized plasmid pBES-004 was introduced into different *B. subtilis* strains (RIK1285, CK7, and 1A1) to study the effect of mannanase expression. However, all exhibited lower activity compared to strain F6004 during PKM fermentation (**Figure 1F**). This may be due to genetic deficiencies affecting cell growth in PKM. Particularly, strain RIK1285 not only has deficiency in major extracellular proteases AprE and NprE, but also mutations affecting tryptophan and lysine biosynthesis, possibly leading to compromised cell growth on PKM where nutrients may not be readily available. In addition, other genetic differences were explored among the strains, focusing on *swrAA*, *sfp*, *bioF*, and *fliF* genes. SwrAA, important for swarming behaviour, is functional in wild-type strains but not in lab strains 1A1 and RIK1285 due to a frameshift mutation (Kamada et al., 2015) **(Supplementary Figure S1A)**. The *sfp* gene, required for producing the antibiotic surfactin, also had a frameshift mutation in 1A1 and RIK1285, preventing surfactin production **(Supplementary Figure S1B)**. None of the strains had deletions in *bioF* or *fliF* genes, which were reported to confer advantages for γ-PGA synthesis (Kamada et al., 2015). The findings emphasize the importance of selecting appropriate host strains for protein expression and specific applications, as genetic variations significantly influence performance.

### 3.2 Improving mannanase production through promoter engineering

Using strong promoter is one of the most efficient methods to increase the production of target proteins. Particularly, transcriptomic data are useful information for screening strong promoters in biotechnological applications. Here, in order to obtain a strong promoter for mannanase expression during PKM fermentation, a transcriptome analysis was conducted to examine the gene expression profile of strain F6 cultivated on solid-state PKM (**Figure 2A**). The highest expression level was found to be from the *ssrA* gene, encoding a hybrid transfer-messenger RNA (tmRNA) molecule involved in the rescue of stalled ribosome during an aberrant translation. Next, among the top 5 expressed genes, there were also genes encoding alkaline serine protease subtilisin (AprE) and neutral metalloprotease (NprE). *B. subtilis* produces eight extracellular proteases to mediate the degradation of extracellular proteins. Among them, AprE and NprE are the most abundant proteases and are found in the culture medium during stationary phase where they contribute more than 95% of the extracellular proteolytic activity of *B. subtilis* (Harwood & Kikuchi, 2022). Given that *aprE* was the second highly expressed gene, this may explain the high mannanase activity driven by *aprE* promoter in the plasmid pBES-004 as observed during PKM fermentation. Other highly expressed genes include *yitM* encoding a biofilm-associated toxin (YIT) which attacks toxin-sensitive competitor cells by passing through the protective barriers of the biofilm. Lastly, there was also *veg* gene which seems to have an important function during the outgrowth of spores (Radeck et al., 2013).

**Figure 2.**
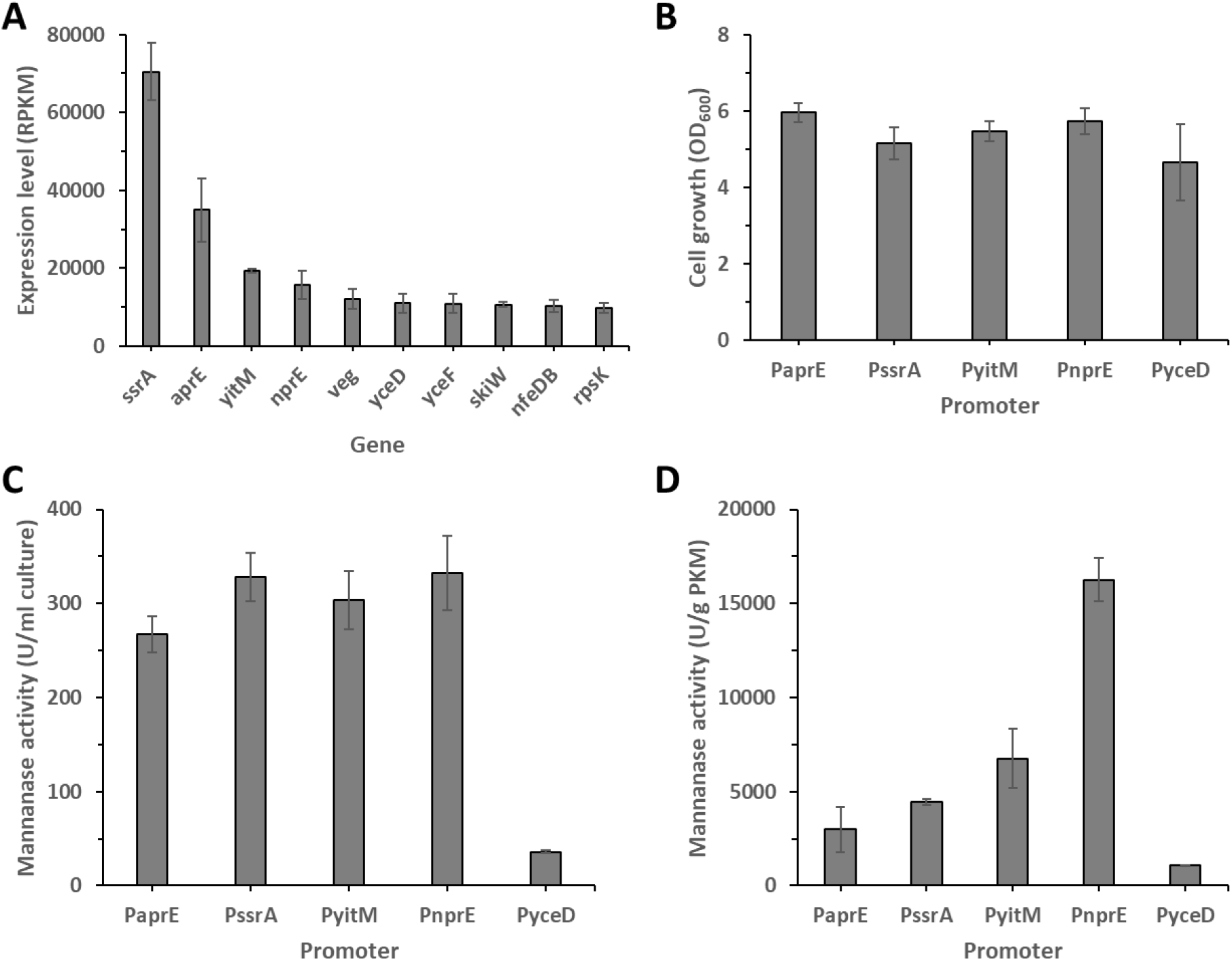
Promoter mining based on transcriptome profile of *B. subtilis* F6. (A) Top 10 highly expressed genes in F6 at the end of 6h-PKM solid-state fermentation. RPKM: Reads Per Kilobase per Million mapped reads. (B, C) (B) Cell growth measurement and (C) mannanase activity of recombinant strains of *B. subtilis* F6 overexpressing mannanase GmuG under different promoters in overnight LB culture. (D) Mannanase activity in PKM solid-state fermentation. Data are the mean of triplicates, and bars indicate standard deviation of triplicates.

We selected five promoters (*aprE, ssrA, yitM, nprE,* and *yceD* promoters) from the highly expressed genes to determine the most suitable one. All of the recombinant strains containing different promoters (F6004, F6004ss, F6004yi, F6004np, F6004yc) exhibited robust cell growth when cultivated in LB medium overnight (**Figure 2B**). In comparison to the control strain F6004 (P*aprE*), the mannanase activity produced by F6004ss (P*ssrA*), F6004yi (P*yitM*), and F6004np (P*nprE*) was slightly higher (**Figure 2C**). In contrast, during PKM fermentation, the mannanase activity produced by F6004np (P*nprE*) was the highest (16263 U/g PKM), which represents an improvement of 5.4-fold compared to the control strain F6004 (P*aprE*) (3007 U/g PKM, **Figure 2D**).

Both proteases AprE and NprE are involved in the supply of amino acids for growth by degradation of extracellular proteins. AprE and NprE may have different physiological functions due to their non-identical regulation and timing of expression. For instance, the expression of *nprE* is regulated by AbrB less tightly than *aprE* expression. It was also observed that each of the promoters jointly repressed by CodY and ScoC displays its own distinct pattern of expression, which likely depends on the relative contributions to regulation and the relative affinities of binding of the two proteins. Therefore, the difference in the regulation of *aprE* and *nprE* expression may account for the remarkable difference in the activity of the respective promoter when used for driving *gmuG* expression during PKM fermentation.

Next, we attempted to further enhance the NprE promoter activity by eliminating the binding sites of the respective repressor. Several NprE promoters with varying lengths (from 82 to 350 bp) were constructed by sequentially excluding the individual transcription factor-binding site (TFBS) (**Figure 3A**). The control strain F6004np contains *nprE* promoter of 250 bp (P*nprE*250) (**Figure 3A**). Firstly, the length of the promoter was extended to 350 bp, forming promoter P*nprE*350 (**Figure 3A**). Recombinant strain F6004np350 (P*nprE*350) exhibited similar level of cell growth and mannanase activity compared to the control strain F6004np (P*nprE*250). This suggests that there is likely no additional TFBSs beyond the 250 bp of promoter sequence. Therefore, P*nprE*250 can be regarded as the intact promoter containing all the native TFBSs. Subsequently, the putative CodY-binding motif was removed from P*nprE*250 and the promoter was shortened to 150 bp, forming promoter P*nprE*150 (**Figure 3A**). Recombinant strain F6004np150 (P*nprE*150) showed a significantly reduced mannanase activity (**Figure 3C, D**) and GmuG expression (**Figure 3E**) despite displaying good cell growth (**Figure 3B**). This could be due to the presence of uncharacterized activator-binding site within the removed sequence. Next, the length of the promoter was further trimmed to 110 bp after removing ScoC-binding site I, generating promoter P*nprE*110 (**Figure 3A**). Recombinant strain F6004np110 (P*nprE*110) produced a remarkably high mannanase activity. The activity was 894 U/ml in LB culture, which is 1.7-fold improvement compared to the control strain F6004np (P*nprE*250) (512 U/ml) (**Figure 3C**). During PKM fermentation, the activity was 20513 U/g PKM, which represents a 1.8-fold increase compared to the control strain F6004np (P*nprE*250) (11153 U/g PKM) (**Figure 3D**). Lastly, the promoter was further shortened to 82 bp after eliminating a large part of the CodY-binding site, yielding promoter P*nprE*82 (**Figure 3A**). However, the mannanase activity produced by the recombinant strain F6004np82 (P*nprE*82) was not significantly different from that of F6004np110 (P*nprE*110). Both the promoters (P*nprE*110 and P*nprE*82) retained the ScoC-binding site II which happened to overlap with the −10 core region.

**Figure 3.**
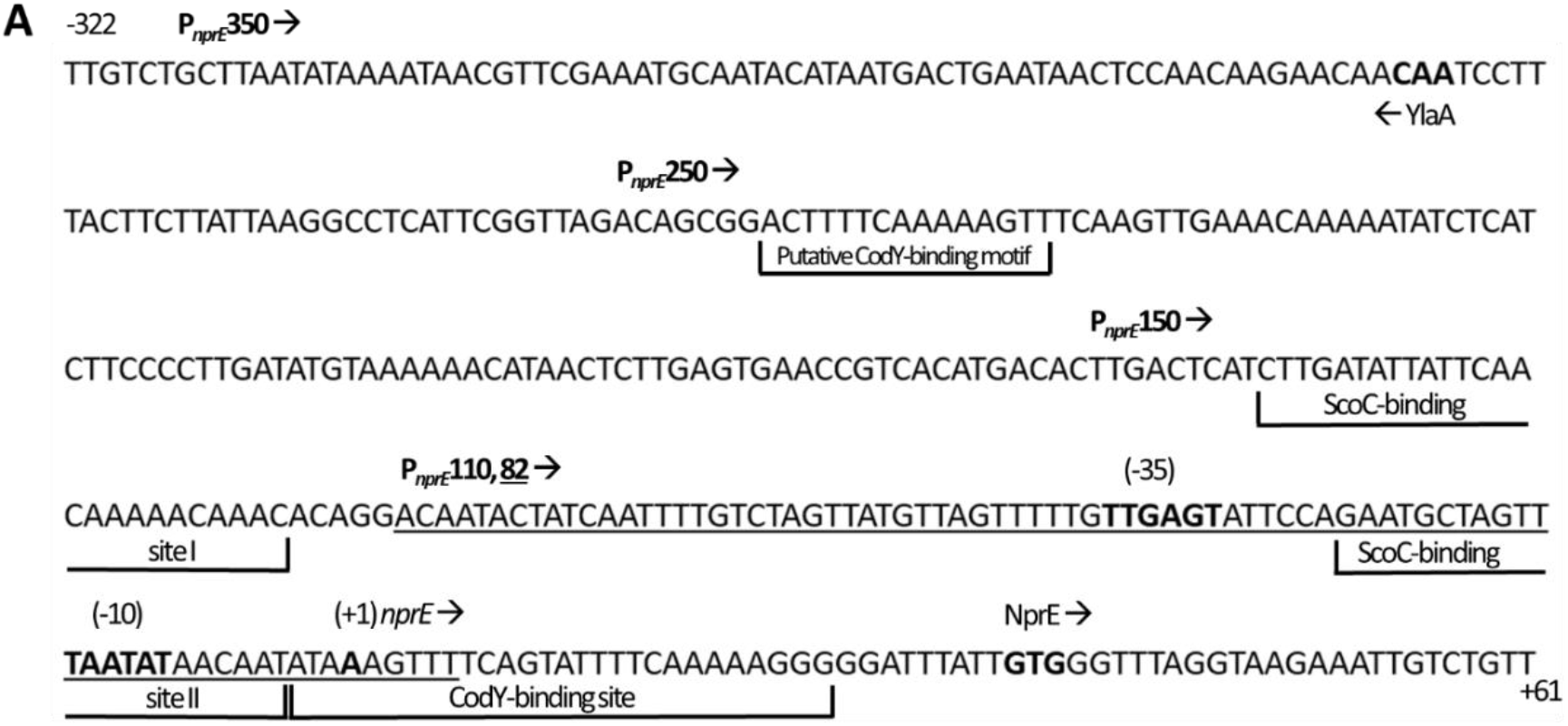
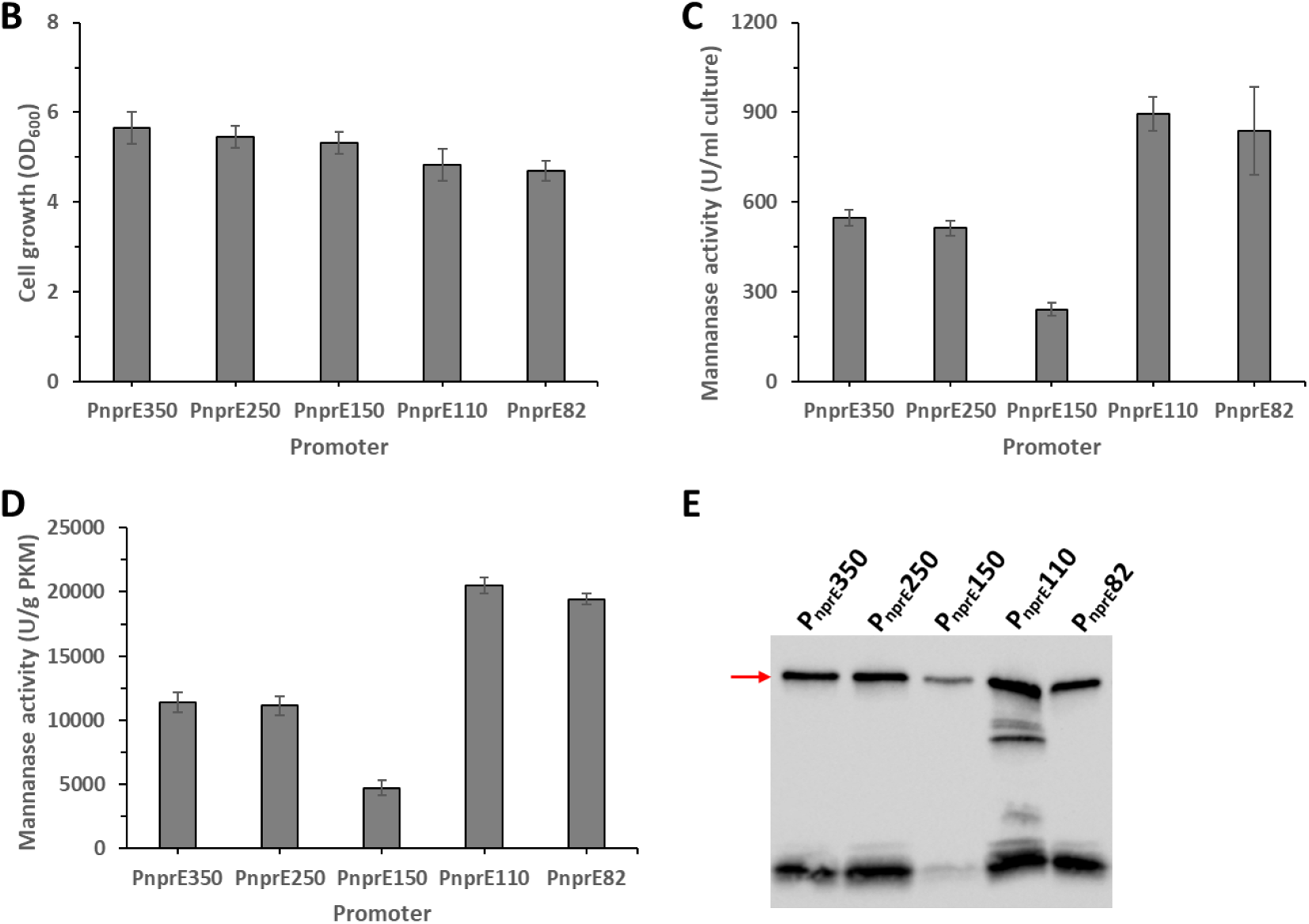
Optimization of *nprE* promoter length. (A) Sequence of the *nprE* regulatory region. Coordinates are reported with respect to the transcription start point. The likely start codon, the −10 and −35 promoter regions, and the transcription start point are shown in bold. The directions of the transcription and translation are indicated by horizontal arrows. The start points of the five different promoters (P*nprE*350, 250, 150, 110, 82) are labelled. All the promoters cover the sequence up to the nucleotide just before NprE start codon, except for P*nprE*82 which has its sequence underlined. (B, C) (B) Cell growth measurement and (C) mannanase activity of recombinant strains of *B. subtilis* F6 overexpressing mannanase GmuG under *nprE* promoter of varying lengths in overnight LB culture. (D, E) (D) Mannanase activity and (E) western blot analysis of GmuG expression in PKM solid-state fermentation. Data are the mean of triplicates, and bars indicate standard deviation of triplicates. The band corresponding to intact GmuG on western blot was marked.

Overall, the highest mannanase activity (**Figure 3C, D**) and protein expression (**Figure 3E**) was observed with F6004np110 (P*nprE*110), where the *nprE* promoter was optimized at 110 bp, containing the core region −35 and −10, ScoC-binding site II, and CodY-binding site (**Figure 3A**). We proposed that the improved promoter activity was mainly attributed to the deletion of ScoC-binding site I in which ScoC may exert a stronger repression on *npre* promoter compared to CodY. It was reported that the repression of *nprE* gene by a derepressed level of ScoC is stronger than that by CodY (Barbieri et al., 2016). However, it is not clear whether this is facilitated by a stronger binding affinity of ScoC to the *nprE* promoter or via other mechanisms. Our findings demonstrated that partial removal of regulatory regions from the NprE promoter could potentially make it become less tightly regulated, resulting in higher gene expression.

### 3.3 Further characterization of F6004np110

A time-course study of PKM solid-state fermentation was performed using the recombinant strains F6004 (P*aprE*), F6004np (P*nprE*250), and F6004np110 (P*nprE*110). As shown in **Figure 4A**, as earlier as 6 h during fermentation, F6004np110 (P*nprE*110) already exhibited significantly higher mannanase activity than the wild-type strain F6. The mannanase activity increased substantially afterward. Overall, compared to the wild-type strain, the engineered strains containing the promoters P*aprE*, P*nprE*250 and P*nprE*110 demonstrated significant enhancements in mannanase activity during fermentation, with improvements of 16-, 25-, and 61-fold, respectively at 30 h (**Figure 4B**). Analysis of protein expression by SDS-PAGE and western blot showed that P*nprE*110 had the highest secretory expression of GmuG protein (**Figure 4C**).

**Figure 4.**
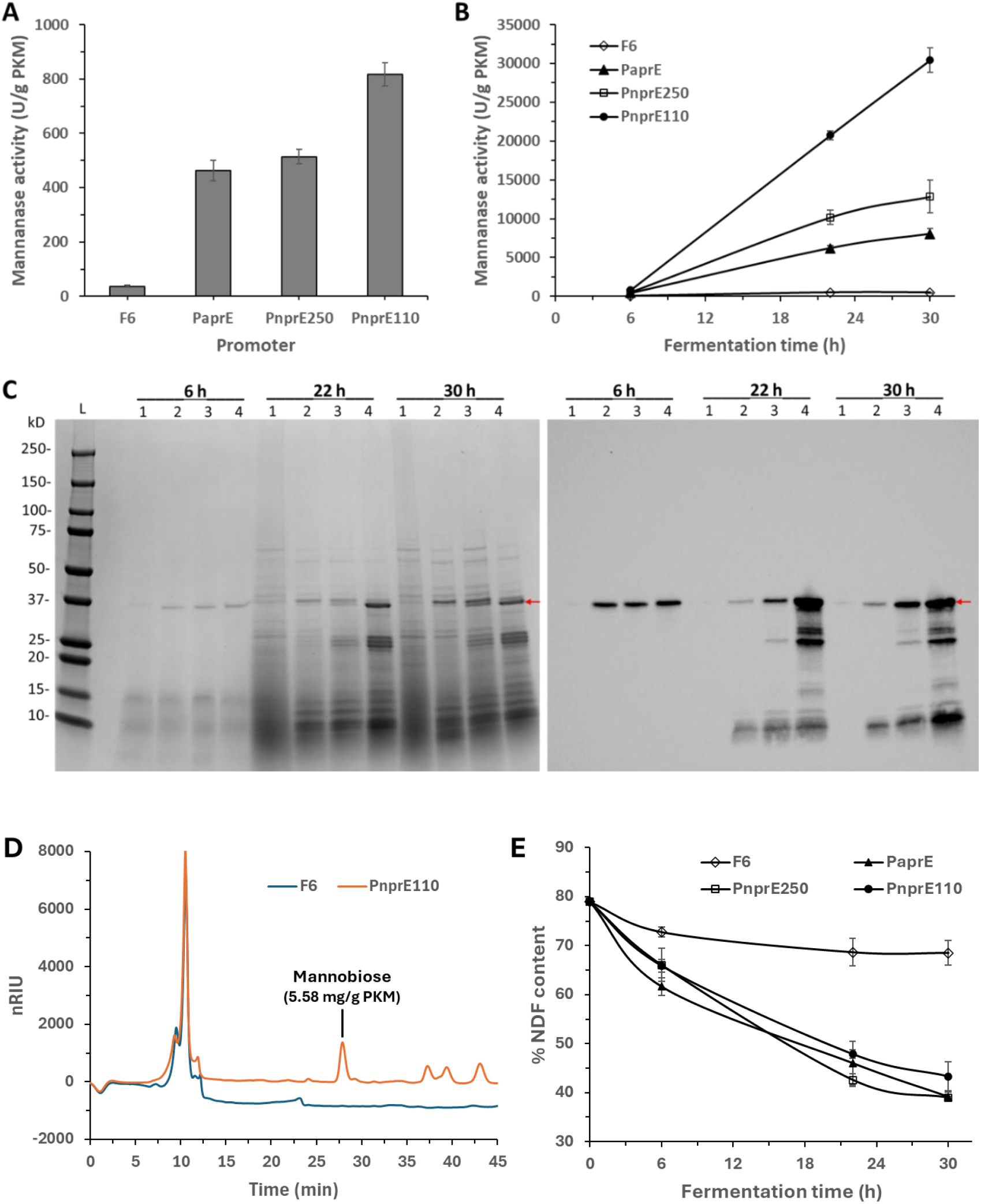
Time-course study of different strains of *B. subtilis* F6 in PKM solid-state fermentation. (A, B) Mannanase activity at (A) 6 h and (B) across different timepoints. (C) SDS-PAGE (left) and western blot (right) analysis of GmuG expression at 6, 22 and 30 h of PKM solid-state fermentation. The band corresponding to intact GmuG was marked. 1: Wild-type F6; 2: F6004 (P*aprE*); 3: F6004np (P*nprE*250); 4: F6004np110 (P*nprE*110). (D) HPLC chromatogram showing the sugar profile of PKM after 30h-fermentation with wild-type F6 or F6004np110 (P*nprE*110). (E) Neutral detergent fibre content of PKM at different timepoints during solid-state fermentation with different F6 strains. Data are the mean of triplicates, and bars indicate standard deviation of triplicates.

Next, the degradation products of PKM fibre hydrolysis were analysed by HPLC. PKM sample was examined for the presence of soluble oligosaccharides or sugar monomers after fermentation with the wild-type strain F6 or the recombinant strain F6004np110 (P*nprE*110). Mannobiose (5.58 mg/g PKM) was detected in F6004np110 (P*nprE*110)-fermented PKM while no oligosaccharide was found in PKM incubated with the wild-type strain F6 (**Figure 4D**). Hence, the presence of mannobiose is indicative of mannan degradation by the recombinant *Bacillus* strain. Our previous work showed that the purified recombinant enzyme of GmuG could release a significant amount of mannobiose and mannotriose from PKM (Ong et al., 2024). Overexpression of GmuG in *Bacillus* likely allowed a higher degree of mannan hydrolysis and accumulation of released sugars during fermentation. PKC-derived mannooligosaccharides (MOS) were found to exhibit prebiotic potential by promoting the growth of probiotics Lactobacilli (Jana & Kango, 2020). Compared to other mannan-rich substrates such as locust bean gum, guar gum, konjac glucomannan and copra meal, the MOS from PKC demonstrated the highest antioxidant, anti-glycating and anti-cancerous activities. Therefore, the generation of beneficial MOS following fermentation using the recombinant strain can potentially enhance the feeding value of PKM as a functional feed ingredient.

Last but not least, the effect of elevated mannanase activity on PKM fibre degradation was assessed using neutral detergent fibre (NDF) analysis. The NDF analysis measures the total fibre in plant cell walls consisting of lignin, cellulose and hemicellulose. On top of this, the NDF value reported in this study also includes the ash content. Before fermentation, the PKM sample showed a high content of NDF fibre at 79% (**Figure 4E**). This is consistent to the measurement performed by Almaguer et al. where the NDF component of PKC was found to be 77.9% (Almaguer et al., 2014). After fermentation with the wild-type strain F6, the fibre content was reduced by 10% in 22 h, with no significant fibre hydrolysis observed afterward. In contrast, the recombinant strain F6004 achieved a 32.9% reduction at 22 h and a 39.8% reduction at 30 h, significantly reducing the NDF content in PKM from 78.9% to 39.2%.

Similarly, after 30 h of fermentation, strains F6004np (P*nprE*250) and F6004np110 (P*nprE*110) exhibited fibre reduction of 40.0% and 35.6%, respectively, comparable to strain F6004. Despite the enhanced mannanase activity achieved through promoter engineering, this did not lead to a greater degree of fibre hydrolysis.

It is assumed that the fermentation process mainly degraded the mannan but not so much other components. Therefore, after excluding 15% lignin and 5% ash from the NDF content, it is estimated that about 20% fibre remained after fermentation. Within the residual fibre, a significant fraction is likely composed of non-mannan polysaccharides based on glucose, xylose, arabinose, and galactose which account for 13.3% of PKC (Cerveró et al., 2010). Microscopy images of PKC cell wall showed an overlapping signal from mannan and cellulose (Gomez-Osorio et al., 2022). The close association between mannan and other polysaccharides may hinder the accessibility of mannan for further hydrolysis. Therefore, to achieve more comprehensive fibre hydrolysis, the supplementation of additional hydrolytic enzymes such as cellulase and xylanase may be beneficial. In addition, the mannanase GmuG may have limited ability to act on the crystalline mannan present in PKM, given that the enzyme has a relatively simple structure without any carbohydrate-binding domains. Hence, it might be beneficial in a future work to explore other heterologous mannanases which may have better capability of breaking down the crystalline mannan in PKM.

### 3.4 Scaled-up solid-state fermentation of PKM

Next, a scaled-up fermentation study was conducted using the highest mannanase-producing strain F6004np110 (P*npre*110). The fermentation involved multi-step blending, with microorganisms added only in the initial step. Then, the pre-fermented PKM and *Bacillus* mixture served as the starter for the following batch. A total amount of 60 g fermented PKM was obtained after 2 days. As shown in **Figure 5A**, the NDF content of PKM was reduced by 29.5% while the protein content was increased slightly by 1.3% after 2 days fermentation. Previous study on PKC fermentation using microbial cocktail of *Bacillus amyloliquefacien* and *Trichoderma harzianum* reported a decrease of NDF content by 6.6% after 7 days incubation (Pasaribu et al., 2019). In comparison, fermentation using the recombinant *B. subtilis* achieved a much higher degree of NDF reduction in a shorter incubation period, highlighting the efficiency of our engineered *bacillus* strain for PKM fibre hydrolysis.

**Figure 5.**
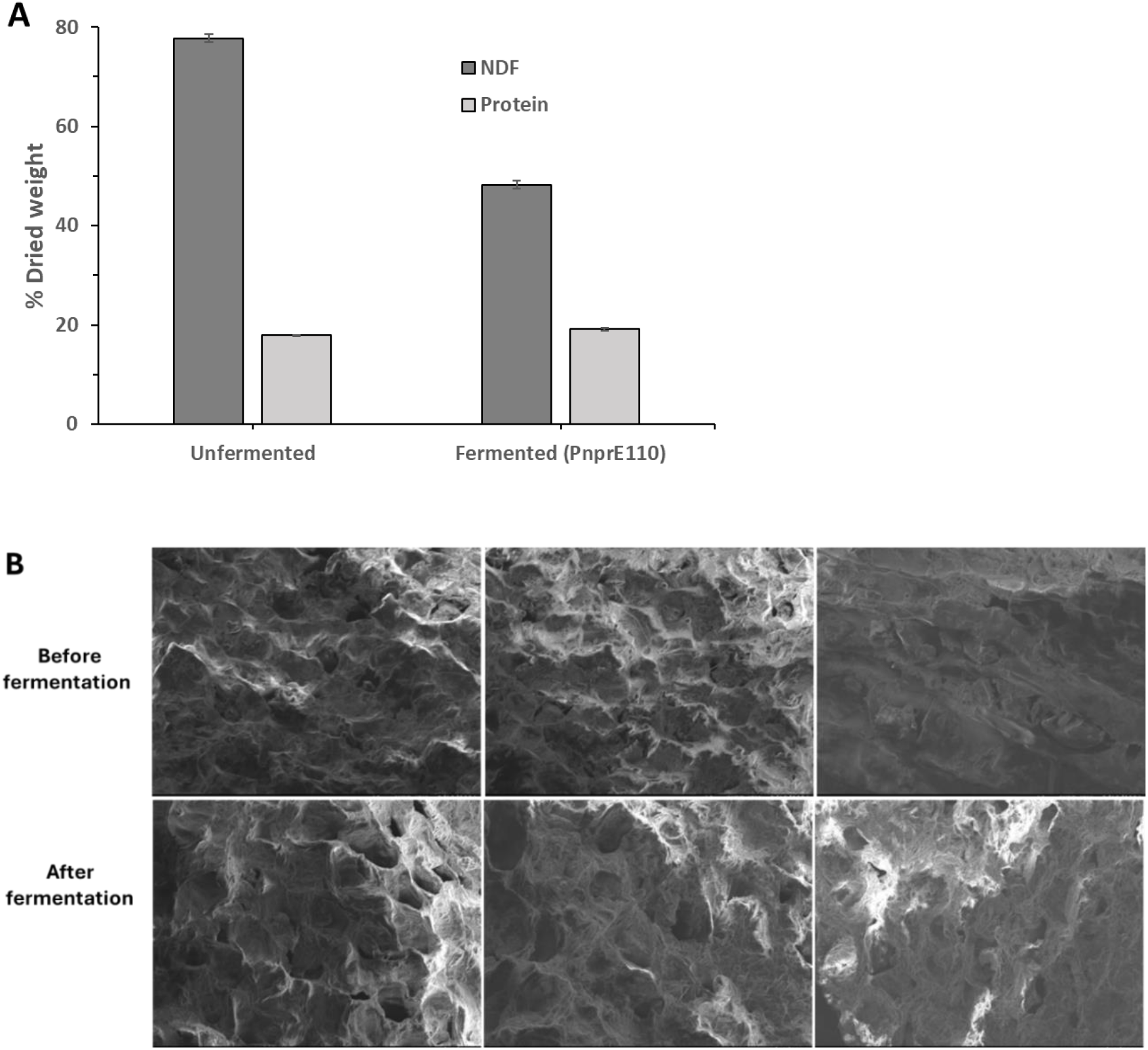
Scaled-up PKM solid-state fermentation using recombinant strain F6004np110 (P*nprE*110). (A) Comparison of neutral detergent fibre (NDF) and protein contents between the unfermented and fermented PKM samples. Data are the mean of triplicates, and bars indicate standard deviation of triplicates. (B) Analysis of surface morphology of PKM under FE-SEM.

Lastly, the surface morphology of PKM was analysed with SEM. After fermentation, the surface of PKM exhibited honeycomb-like structures, and appeared more porous (**Figure 5B**). This is consistent with the observation by Wang et al. where the surface of millet bran displayed honeycomb-like structures and became loose and porous after solid-state fermentation with cellulase-producing microbes and cellulase supplementation (Wang et al., 2023). The increased porosity on the PKM surface could be attributed to the enzymatic hydrolysis of mannan and the disruption of the cell wall structure during the microbial fermentation. Overall, our study suggests that the optimized recombinant strain F6004np110 (P*npre*110) can be applied for scaled-up PKM fermentation, resulting in significant reduction in NDF content. Further optimization of the fermentation conditions (e.g., aeration, mixing) may be helpful in improving the efficiency of fibre hydrolysis in large scale.

## 4. Conclusion

In this study, the mannanase production in the undomesticated strain of *B. subtilis* F6 was enhanced by homologous expression of mannanase GmuG using a replicative plasmid. Plasmid optimization was performed systematically to select the best combination of RBS, SP and promoter for high mannanase expression. The highest mannanase activity was obtained from the recombinant strain F6004np110 (30425.4 U/ g PKM), representing a remarkable 61- fold increase compared to the native strain F6 after 30 h of PKM fermentation. While the native strain F6 reduced the NDF content by 10%, the recombinant strains achieved a substantial 35 - 40% decrease in the NDF content. Our study showed that protein expression can be enhanced by optimizing RBS and SP in which their interactions may potentially affect the mRNA secondary structure around the translation initiation site, thereby impacting the efficiency of translation initiation. Furthermore, a significant increase in mannanase activity was obtained through promoter mining based on the genes highly expressed during PKM fermentation, highlighting the efficacy of transcriptome data in identifying strong promoters under specific growth conditions.

Given the potential strain-to-strain variations in adaptation to solid-state PKM, selecting an optimal host strain for genetic engineering is crucial to ensure the performance of the recombinant strain in intended applications. Environmental isolates sharing common features like swarming and surfactin production may offer advantages over laboratory strains when cultivating on unconventional substrates. Given the concerns raised about the potential transfer of antibiotic resistance genes from genetically modified organisms to pathogenic bacteria in the environment, future work can focus on developing antibiotic-free selection which avoids the use of antibiotic resistance genes as selection markers. An alternative selection method is essential gene complementation. However, this method requires engineering mutant host strains and using specifically defined media. Overall, the study demonstrates the construction of a high mannanase expression system in an undomesticated strain of *B. subtilis* through systematic plasmid optimization. The high mannanase expression was proven to significantly improve NDF degradation during solid-state fermentation of PKM, and this could potentially enhance the digestibility and feeding value of PKM for non-ruminant livestock.

## List of abbreviations

DNS: Dinitrosalicylic acid
EDTA: Disodium ethylene diamine tetraacetate
FE-SEM: Field emission scanning electron microscopy
GRAS: Generally regarded as safe
HPLC: High-performance liquid chromatography
LB: Lysogeny broth
LBG: Locust bean gum
MBP: Maltose binding protein
MOS: Mannooligosaccharide
NDF: Neutral detergent fibre
nRIU: Nano refractive index units
P: Promoter
PBS: Phosphate-buffered saline
PKC: Palm kernel cake
PKE: Palm kernel expeller
PKM: Palm kernel meal
RBS: Ribosome binding site
RID: Refractive index detector
RIN: RNA integrity number
RPKM: Reads per kilobase of transcript per million mapped reads
SDS-PAGE: Sodium dodecyl sulfate polyacrylamide gel electrophoresis
SP: Signal peptide
SSF: Solid-state fermentation
TFBS: Transcription factor-binding site
tmRNA: Transfer-messenger RNA

## Acknowledgements

The authors would like to thank Professor Chua Nam Hai for his constructive comments on this project, and Mr. Hermil Calasang for arrangement of PKM sample.

