## Supplemental Figure S1 for "Improving Mannanase Production in *Bacillus subtilis* for Fibre Hydrolysis during Solid-State Fermentation of Palm Kernel Meal"

**Supplementary figure:**

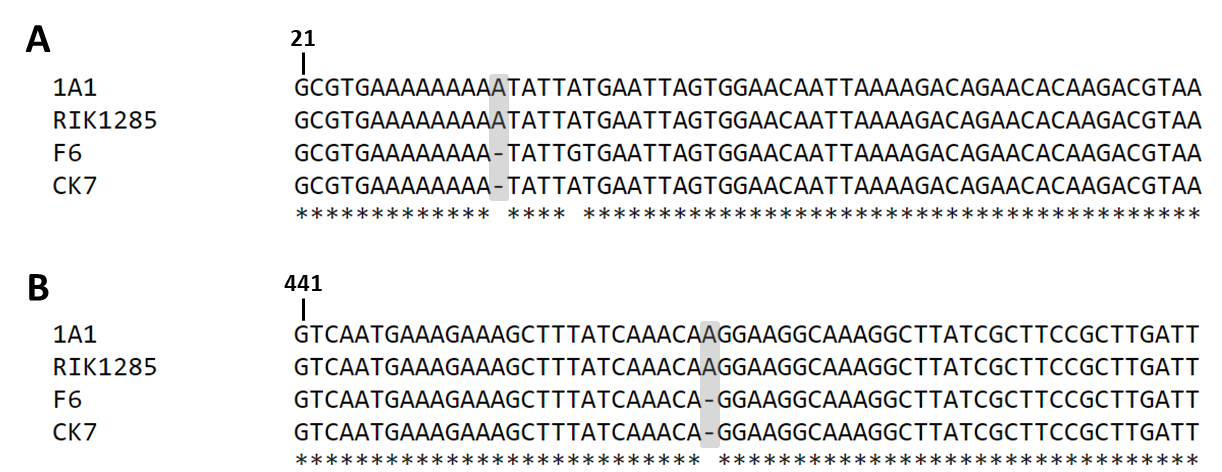

**Figure S1** **Genetic differences among different *B. subtilis* strains (1A1, RIK1285, F6 and CK7).** Sequence alignment of selected region of (A) *swrAA* and (B) *sfp* genes. The highlighted region shows the adenine insertion occurred in strains 1A1 and RIK1285.
